# immuneML: an ecosystem for machine learning analysis of adaptive immune receptor repertoires

**DOI:** 10.1101/2021.03.08.433891

**Authors:** Milena Pavlović, Lonneke Scheffer, Keshav Motwani, Chakravarthi Kanduri, Radmila Kompova, Nikolay Vazov, Knut Waagan, Fabian L. M. Bernal, Alexandre Almeida Costa, Brian Corrie, Rahmad Akbar, Ghadi S. Al Hajj, Gabriel Balaban, Todd M. Brusko, Maria Chernigovskaya, Scott Christley, Lindsay G. Cowell, Robert Frank, Ivar Grytten, Sveinung Gundersen, Ingrid Hobæk Haff, Sepp Hochreiter, Eivind Hovig, Ping-Han Hsieh, Günter Klambauer, Marieke L. Kuijjer, Christin Lund-Andersen, Antonio Martini, Thomas Minotto, Johan Pensar, Knut Rand, Enrico Riccardi, Philippe A. Robert, Artur Rocha, Andrei Slabodkin, Igor Snapkov, Ludvig M. Sollid, Dmytro Titov, Cédric R. Weber, Michael Widrich, Gur Yaari, Victor Greiff, Geir Kjetil Sandve

## Abstract

Adaptive immune receptor repertoires (AIRR) are key targets for biomedical research as they record past and ongoing adaptive immune responses. The capacity of machine learning (ML) to identify complex discriminative sequence patterns renders it an ideal approach for AIRR-based diagnostic and therapeutic discovery. To date, widespread adoption of AIRR ML has been inhibited by a lack of reproducibility, transparency, and interoperability. immuneML (immuneml.uio.no) addresses these concerns by implementing each step of the AIRR ML process in an extensible, open-source software ecosystem that is based on fully specified and shareable workflows. To facilitate widespread user adoption, immuneML is available as a command-line tool and through an intuitive Galaxy web interface, and extensive documentation of workflows is provided. We demonstrate the broad applicability of immuneML by (i) reproducing a large-scale study on immune state prediction, (ii) developing, integrating, and applying a novel method for antigen specificity prediction, and (iii) showcasing streamlined interpretability-focused benchmarking of AIRR ML.

## Introduction

T-cell receptors (TCRs) and B-cell receptors (BCRs), that are collectively known as adaptive immune receptor (AIR) repertoires (AIRRs), recognize antigens and record information on past and ongoing immune responses^1–4^. AIRR-encoded information is particularly useful for the *repertoire*-based prediction and analysis of immune states (e.g., health, disease, infection, vaccination) in relation to other metadata such as major histocompatibility complex (MHC)^5–7^, age^7,8^, and sex^9^. Together this information shapes the foundation for AIRR-based diagnostics^6,10–14^. Similarly, *sequence*-based prediction of antigen and epitope binding is of fundamental importance for AIR-based therapeutics discovery and engineering^15–25^. In this manuscript, the term *AIRR* signifies both AIRs and AIRRs (a collection of AIRs) if not specified otherwise.

Machine learning (ML) has recently entered center stage in the biological sciences because it allows detection, recovery, and re-creation of high-complexity biological information from large-scale biological data^26–29^. AIRRs have complex biology with specialized research questions, such as immune state and receptor specificity prediction, that warrant domain-specific ML analysis^15^. Briefly, (i) ~10^8^–10^10^ distinct AIRs exist in a given individual at any one time^30–32^, with little overlap among individuals, necessitating encodings that allow detection of predictive patterns. These shared patterns may correspond to full-length AIRs^6^ or subsequences^16^ alternative representations thereof^11,12,17,18,22,33–35^. (ii) In repertoire-based ML, the patterns relevant to any immune state may be as rare as one antigen-binding AIR per million lymphocytes in a repertoire^36^ translating into a very low rate of relevant sequences per repertoire (low witness rate)^11,37,38^. (iii) In sequence-based ML, the enormous diversity of antigen recognition combined with polyreactivity points to complex high-order statistical dependencies in the short sequence known to be the main determinant of antigen recognition (complementarity-determining region 3, CDR3)^1,16^.

Tailored ML frameworks and platforms that account for the idiosyncrasies of the underlying data have been published for applications in genomics^39,40^, proteomics^41,42^, biomedicine^43^, and chemistry^44^. Their creation recognizes the infeasibility to define, implement, and train appropriate ML models by relying solely on generic ML frameworks such as scikit-learn^45^ or PyTorch^46^. The lack of a standardized framework for AIRR ML has led to heterogeneity in terms of technical solutions, domain assumptions, and user interaction options, hampering transparent comparative evaluation and the ability to explore and select the ML methodology most appropriate for a given study^15^.

## Results

### immuneML overview

Here, we present immuneML, an open-source collaborative ecosystem for AIRR ML (Figure 1). immuneML enables the ML study of both experimental and synthetic AIRR-seq data that are labeled on the repertoire-level (e.g., immune state, sex, age, or any other metadata) or sequence-level (e.g., antigen binding), all the way from preprocessing to model training and model interpretation. It natively implements model selection and assessment procedures like nested cross-validation to ensure robustness in selecting the ML model. immuneML may be operated either via the command line or the Galaxy web interface^47^, which offers an intuitive user interface that promotes collaboration and reusability through shareable analysis histories. To expedite analyses, immuneML may also be deployed to cloud services such as Amazon Web Services (AWS) and Google Cloud, or on a local server for data privacy concerns. Reproducibility and transparency are achieved by shareable specification files, which include all analysis details (Supplementary Figure 1). immuneML’s compliance with AIRR community software and sequence annotation standards^48,49^ ensures straightforward integration with third-party tools for AIRR data preprocessing and AIRR ML results’ downstream analysis. For example, immuneML is fully compatible with the sequencing read processing and annotation suite MiXCR^50^ and the Immcantation^51,52^ and immunarch^53^ frameworks for AIRR data analysis. AIRR data from the AIRR Data Commons^54^ through the iReceptor Gateway^55^, as well as the epitope-specific TCR database VDJdb^56^ may be directly downloaded into the immuneML Galaxy environment. Additionally, immuneML is integrated with the AIRR-specific attention-based multiple-instance learning ML method DeepRC^37^, the TCR-specific clustering method TCRdist^17^, and is compatible with GLIPH2^57^.

**Figure 1.**
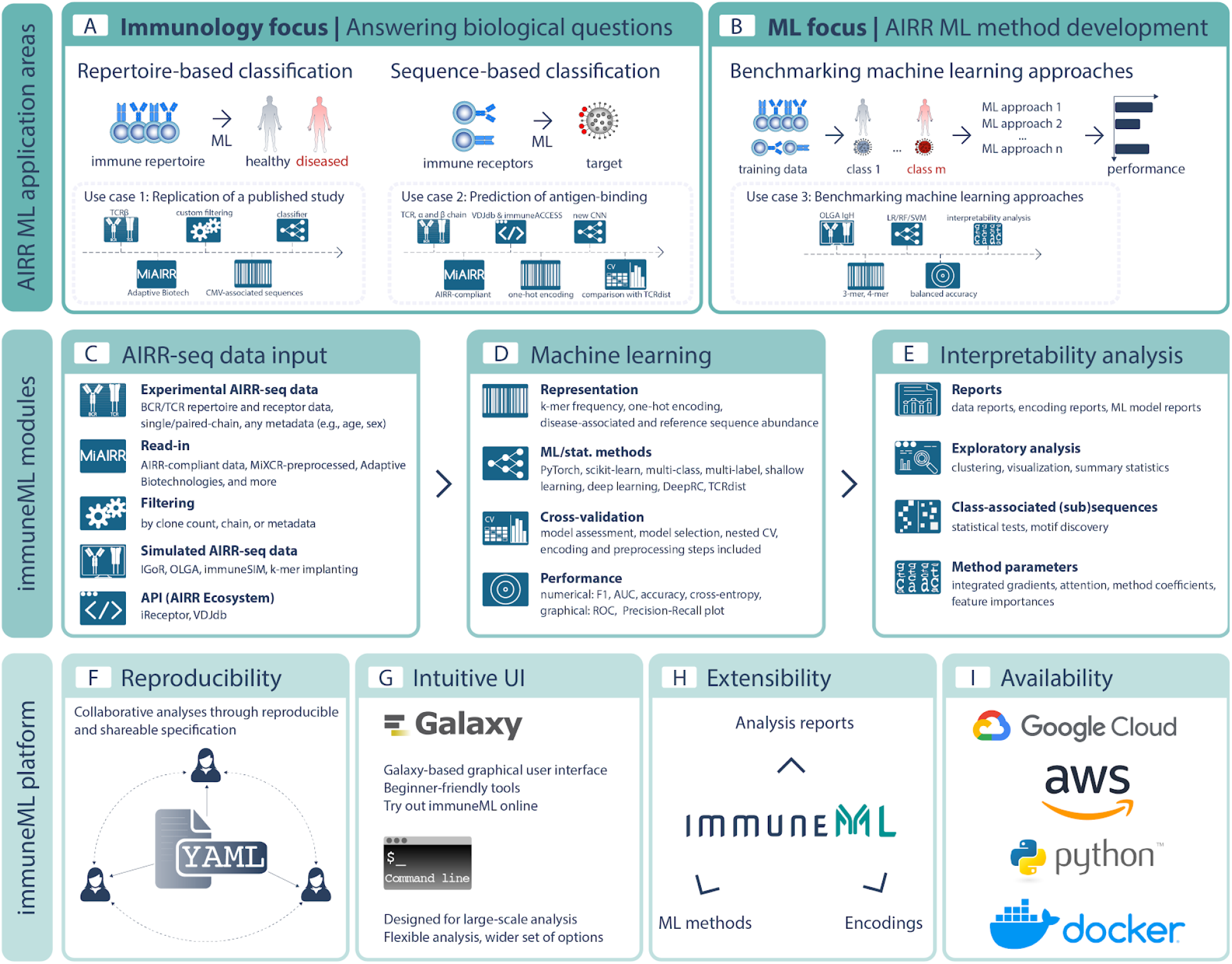
Overview of immuneML. (**A**) The main immuneML application areas are sequence- and repertoire-based prediction of AIRR with application to (**A**) immunodiagnostics and therapeutics research, as well as to (**B**) develop AIRR-based methods. We show three use cases belonging to these application areas. <u>Use case 1</u>: reproduction of the study by Emerson et al.^6^ on repertoire classification, <u>use case 2</u>: extending the platform with a novel convolutional neural network (CNN) classifier for prediction of TCR-pMHC binding that allows paired-chain input, <u>use case 3</u>: benchmarking ML methods with respect to their ability to recover a sequence-implanted immune signal. The immuneML core is composed of three pillars, which are (**C**) AIRR-seq data input and filtering, (**D**) ML, and (**E**) Interpretability analysis. Each of these pillars has different modules that may be interconnected to build an immuneML workflow. (**F**) immuneML uses a specification file (YAML), which is customizable and allows full reproducibility and shareability with collaborators or the broader research community. An overview of how immuneML analyses can be specified is given in Supplementary Figure 1. **(G)** immuneML may be operated via the Galaxy web interface or the command line. (**H**) All immuneML modules are extendable. Documentation for developers is available online. (**I**) immuneML is available as a Python package, a Docker image, and may be deployed to cloud frameworks (e.g., AWS, Google Cloud). Abbreviations: CMV (cytomegalovirus).

To get started with immuneML, we refer the reader to **Focus Box 1**. To demonstrate immuneML’s capabilities for performing AIRR ML, we provide an overview of the main features of the platform, and then highlight three orthogonal use cases: (i) we reproduce the cytomegalovirus (CMV) serostatus prediction study of Emerson et al.^6^ inside immuneML and examine the robustness of the approach showing one way of using immuneML for repertoire-based immune state prediction, (ii) we apply a new custom convolutional neural network (CNN) for the sequence-based task of antigen-binding prediction based on paired-chain TCR data and (iii) we show the use of immuneML for benchmarking AIRR ML methods.

### immuneML allows read-in of experimental single- and paired-chain data from offline and online sources as well as the generation of synthetic data for ML benchmarking

Experimental data may be read-in directly if it complies with the major formats used for AIRR-seq data V(D)J annotation: AIRR-C standard-conforming^48^, MIXCR^50^, 10x Genomics^58^, Adaptive Biotechnologies ImmunoSEQ^6,59^ or VDJdb formats^56^. The AIRR-C format compatibility ensures that also synthetic data as generated by immuneSIM^60^ can be imported. Importing synthetic data as generated by IGoR^61^ and OLGA^62^ is also supported. Moreover, immuneML can be configured to read in data from any custom tabular format. To facilitate access to large-scale AIRR-seq data repositories, we provide Galaxy^47^ tools to download data from the AIRR Data Commons^54^ via the iReceptor Gateway^55^ and from VDJdb^56^ into the Galaxy environment for subsequent ML analysis. Furthermore, immuneML includes built-in capacities for complex synthetic AIRR data generation to satisfy the need for ground-truth data in the context of ML method benchmarking. Finally, read-in data may be filtered by clone count, metadata, and chain.

### immuneML supports multiple ML frameworks and allows for interpretation of ML models

immuneML supports two major ML platforms to ensure flexibility: scikit-learn^45^ and PyTorch^46^ and, therefore, is compliant with all ML methods inside these platforms. immuneML features scikit-learn implementations such as logistic regression, support vector machine, and random forest. In addition, we provide AIRR-adapted ML methods. Specifically, for repertoire classification, immuneML includes a custom implementation of the method published by Emerson et al.^6^, as well as the attention-based deep learning method DeepRC^37^. For paired-chain sequence-based prediction, immuneML includes a custom-implemented CNN-based deep learning method, integrates with TCRdist^17^, and is compatible with GLIPH2^57^. immuneML also includes several encodings that are commonly used for AIRR data such as k-mer frequency decomposition, one-hot encoding where each position in the sequence is represented by a vector of zeros except one entry containing 1 denoting appropriate amino acid or nucleotide, encodings by the presence of disease-associated sequences, and repertoire distances. For the full overview of analysis components, see Supplementary Table 1.

A variety of tabular and graphical analysis reports may be automatically generated as part of an analysis, providing details about the encoded data (e.g., feature value distributions), the ML model (e.g., interpretability reports), and the prediction accuracy (a variety of performance metrics across training, validation, and test sets). Additionally, the trained models may be exported and used in future analyses.

### immuneML facilitates reproducibility, interoperability, and transparency of ML models

immuneML draws on a broad range of techniques and design choices to ensure that it meets the latest expectations with regard to usability, reproducibility, interoperability, extensibility, and transparency^63–66^ (Figure 1).

#### Usability

is achieved by a range of installation and usage options, catered to novices and experts, and to small and large-scale analyses. A Galaxy web interface^47^ allows users to run analyses without the need for any installation and without requiring any skills in programming or command-line operations. Availability through GitHub, pip, and Docker streamlines usage at scales ranging from laptops to high-performance infrastructures such as Google Cloud and AWS (docs.immuneml.uio.no/installation/cloud.html).

#### Reproducibility

is ensured by leveraging the Galaxy framework^47^ that enables sharing of users’ analysis histories, including the data and parameters, so that they can be independently reproduced. If working outside Galaxy, reproducibility is ensured by shareable analysis specification (YAML) files. YAML specification files produced in the Galaxy web interface can also be downloaded to seamlessly switch between Galaxy and command-line operation.

#### Interoperability

is ensured by supporting the import from multiple data sources and export into AIRR-C format (MiAIRR standard) for post-analysis by third-party tools that are AIRR-compliant^48^.

#### Extensibility

of immuneML, signifying straightforward inclusion of new ML methods, encodings, reports, and preprocessing, is ensured by its modular design (Supplementary Figure 2). The code is open-source and available on GitHub (**Focus Box 2**). The documentation details step-by-step developer tutorials for immuneML extension (docs.immuneml.uio.no/developer_docs.html).

#### Transparency

is established by (i) a YAML analysis specification in which the assumptions of the AIRR ML analysis are explicitly defined, and default parameter settings are exported, (ii) separate immunologist-centric Galaxy user interfaces that translate parameters and assumptions of the ML process to aspects of immune receptors that immunologists may better relate to (Supplementary Figure 3) and (iii) for each analysis report, the availability of underlying data for further user inspection.

##### Focus Box 1: Getting started with immuneML

➔ Visit the project website at immuneml.uio.no. immuneML may be used (i) online via the Galaxy web interface (galaxy.immuneml.uio.no), (ii) through a Docker container, or (iii) from the command line by installing and running immuneML as a Python package. Detailed instructions for each of these options are available in the immuneML documentation: docs.immuneml.uio.no/installation.html.

###### Getting started: web interface

➔ For immunologists, we recommend the Quickstart guide based on simplified interfaces for training ML models: docs.immuneml.uio.no/quickstart/galaxy_simple.html. Explanations of the relevant ML concepts can be found in the documentation (sequence classification docs.immuneml.uio.no/galaxy/galaxy_simple_receptors.html and repertoire classification docs.immuneml.uio.no/galaxy/galaxy_simple_repertoires.html)
➔ Alternatively, to have full control over all details of the analysis, see the YAML-based Galaxy Quickstart guide: docs.immuneml.uio.no/quickstart/galaxy_yaml.html.
➔ For guidance on how to use each immuneML Galaxy tool, see the immuneML & Galaxy documentation (docs.immuneml.uio.no/galaxy.html) and the list of published example Galaxy histories (galaxy.immuneml.uio.no/histories/list_published).

###### Getting started: command-line interface

➔ For the command-line Quickstart guide, see docs.immuneml.uio.no/quickstart/cli_yaml.html
➔ For detailed examples of analyses that can be performed with immuneML, see the tutorials (docs.immuneml.uio.no/tutorials.html), use case examples (docs.immuneml.uio.no/usecases.html), and see all supported analysis options in the YAML specification documentation (docs.immuneml.uio.no/specification.html). For any questions, contact us at, visit the troubleshooting page in the documentation (docs.immuneml.uio.no/troubleshooting.html), or open an issue on our GitHub repository (github.com/uio-bmi/immuneML/issues).

##### Focus Box 2: How to contribute to immuneML

There exist multiple avenues for contributing and extending immuneML:

➔ ML workflows for specific research questions can be shared on galaxy.immuneml.uio.no, which allows other researchers to use them directly in their own data analysis.
➔ Questions, enhancements, or encountered bugs may be reported on the immuneML GitHub under “Issues” (github.com/uio-bmi/immuneML/issues).
➔ To improve or extend the immuneML platform, obtain the source code from GitHub at github.com/uio-bmi/immuneML. The immuneML codebase is described in the immuneML developer documentation docs.immuneml.uio.no/developer_docs.html, along with tutorials on how to add new ML methods, encodings, and report components to the platform.
➔ We encourage developers to contribute their improvements and extensions back to the community, either by making their own versions public or by submitting their contributions as GitHub “pull requests” to the main immuneML codebase.

**Figure 2.**
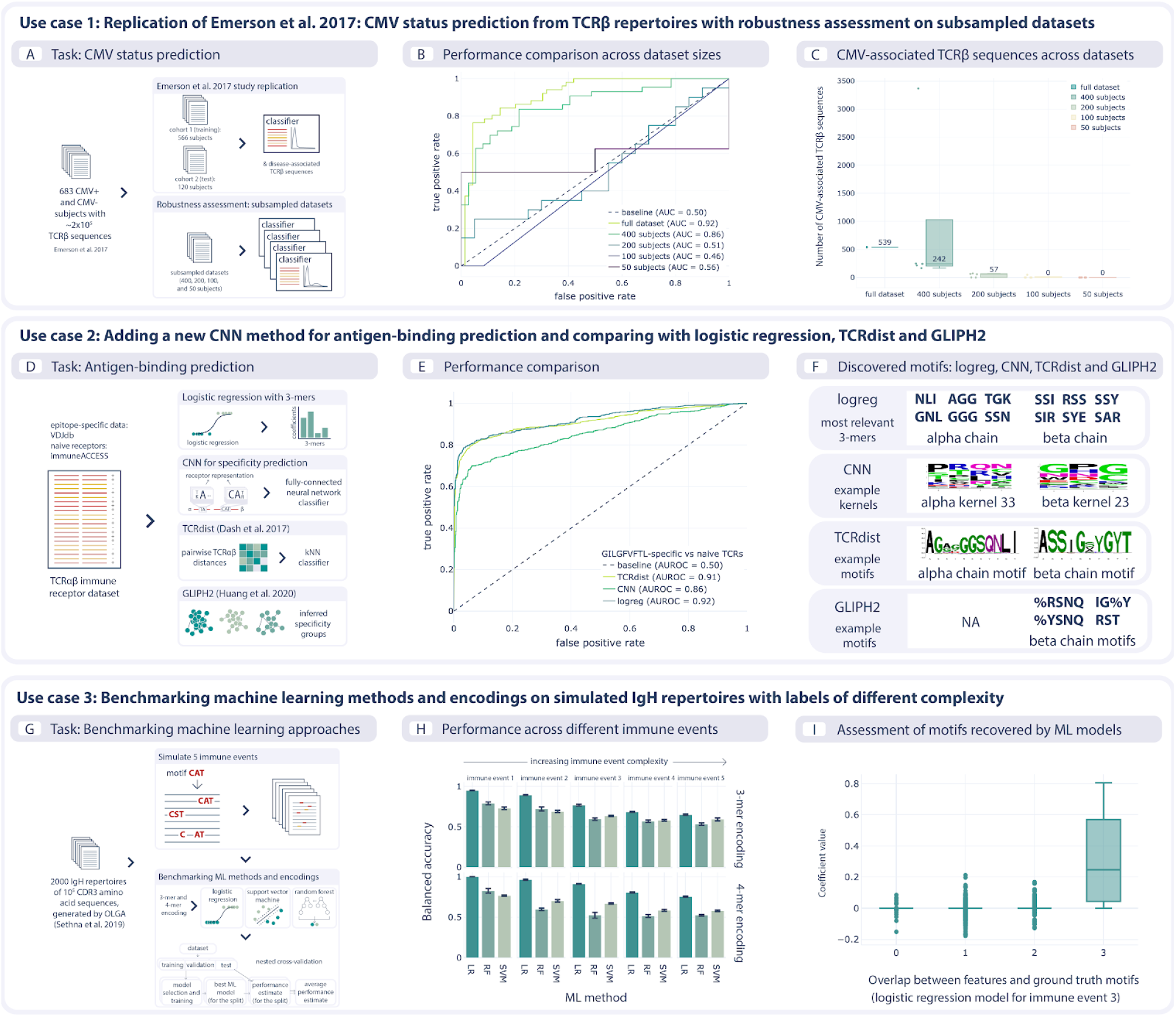
Use cases demonstrating ML model training, benchmarking, and platform extension. We showcase three use cases to exemplify immuneML usage. **(A–C)** <u>Use case 1</u>: Reproduction of a published study^6^ where the task consisted in distinguishing between TCRβ repertoires from CMV (cytomegalovirus) positive and negative individuals, as well as the identification of TCRβ sequences that are associated with CMV status. In addition, we assessed the robustness of the respective statistical approach, measured by the predictive performance, as a function of decreasing dataset size. We show how a lower number of repertoires (400, 200, 100, and 50) leads to decreased prediction accuracy (AUROC: 0.86–0.46) and a lower number of CMV-associated TCRβ sequences (with almost none found in datasets of 100 and 50 subjects). **(D–F)** <u>Use case 2</u>: We developed a new ML method for antigen-specificity prediction on paired-chain T-cell receptor data using a convolutional neural network (CNN) architecture. The method separately detects motifs in paired chains and combines the motif scores corresponding to kernel activations to obtain the receptor representation which is then used as input to a classifier. We compared the CNN method with the TCRdist-based k-nearest neighbor classifier and logistic regression on a dataset consisting of epitope-specific and naive TCRαβ sequences (assumed to be non-epitope-specific). For epitope-specific sequences, we used Epstein-Barr-virus-specific TCRαβ sequences binding to the GILGFVFTL epitope. We also show the motifs recovered by CNN, TCRdist, and GLIPH2 among the epitope-specific sequences. **(G–I)** <u>Use case 3</u>: We show how ground-truth synthetic data may be used to benchmark AIRR ML methods. The dataset consists of 2000 immune repertoires generated by OLGA^62^. Using immuneML, five immune events of increasing complexity are simulated by implanting synthetic signals into the OLGA-generated repertoires. This dataset is subsequently used to benchmark three different ML methods (logistic regression (LR), support vector machine (SVM), and random forest (RF)) in combination with two encodings (3-mer and 4-mer encoding) inside immuneML, showing that the classification performance drops as the immune event complexity increases. The quality of the ML models was further assessed by comparing the feature coefficient sizes with how well these features represent the ground-truth signals. This revealed that models with a good classification performance were indeed able to recover the ground-truth signals.

### Use case 1: Reproduction of a published study inside immuneML

To show how a typical AIRR ML analysis may be performed within immuneML, we reproduced a previously published study by Emerson et al. on the TCRβ-repertoire-based classification of individuals into CMV seropositive and seronegative^6^ (Figure 2A). Using the standard interface of immuneML, we set up a repertoire classification analysis using 10-fold cross-validation on cohort 1 of 563 patients to choose optimal hyperparameters for immuneML’s native implementation of the statistical classifier introduced by Emerson and colleagues. We then retrained the classifier on the complete cohort 1 and tested it on a second cohort (cohort 2) of 120 patients, as described in the original publication (see Methods).

immuneML exports classifier details, such as a list of immune-status-associated sequences for each classifier created during cross-validation, as well as a performance overview using the metrics of choice. We replicated the predictive performance achieved by Emerson et al.^6^, finding 143 of the same CMV-associated TCRs (out of 164) reported in the original study.

We further used built-in robustness analysis of immuneML to explore how classification accuracy and the set of immune-status-associated sequences varied when learning classifiers based on smaller subsets of repertoires (Figure 2 A and B). While the exact set of learned immune-status-associated sequences varied across subsampled data of sizes close to the full dataset, the classification accuracy was nonetheless consistently high (>0.85) as long as the number of training repertoires was 400 or higher (below this, classification accuracy on the separate test sets deteriorated sharply) (Figure 2 B and C). The YAML specification files and detailed results are provided in the immuneML use case documentation (docs.immuneml.uio.no/usecases/emerson_reproduction.html).

### Use case 2: Extending immuneML with a deep learning component for antigen specificity prediction based on paired-chain (single immune cell) data

To illustrate the extensibility of the immuneML platform, we added a new CNN component for predicting antigen specificity based on paired-chain AIR data. The ML task is to discover motifs in the two receptor chains (sequences) and to exploit the presence of these motifs to predict if the receptor will bind the antigen. As the immuneML platform provides comprehensive functionality for parsing and encoding paired-chain data, for hyperparameter optimization, and for presenting results, the only development step needed was to add the code for the CNN-based method itself (Supplementary Figure 5). Briefly, the added CNN consists of a set of kernels for each chain that act as motif detectors, a vector representation of the receptor obtained by combining all kernel activations, and a fully-connected layer that predicts if the receptor will bind the antigen or not. Furthermore, we show how to run analyses with the added component and compare its results with those of alternative models, such as a logistic regression model based on 3-mer frequencies and a k-nearest neighbor classifier relying on TCRdist^17^ as the distance metric (available directly from immuneML through the tcrdist3 package^67^). We also show that the motifs can be recovered from the CNN model, the logistic regression, TCRdist, and GLIPH2^57^ (Figure 2 D).

### Use case 3: ML methods benchmarking on ground-truth synthetic data

Given the current rise in AIRR ML applications, the ability for method developers and practitioners to efficiently benchmark the variety of available approaches is becoming crucial^1,15,60^. Due to the limited current availability of high-resolution, labeled experimental data, rigorous benchmarking relies on a combination of experimental and simulated ground-truth data. The immuneML platform natively supports both the generation of synthetic data for benchmarking purposes and the efficient comparative benchmarking of multiple methodologies based on synthetic as well as experimental data. To exhibit the efficiency with which such benchmarking can be performed within the immuneML framework, we simulated, using the OLGA framework^62^, 2000 human IgH repertoires consisting of 10^5^ CDR3 amino acid sequences each, and implanted sequence motifs reflecting five different immune events of varying complexity (Figure 2 G, Supplementary Table 2). We examined the classification accuracy of three assessed ML methods (Figure 2 H) and used a native immuneML report to examine the overlap between ground truth implanted motifs and learned model features (Figure 2 I, Supplementary Figure 6).

## Discussion

We have presented immuneML, a collaborative and open-source platform for transparent AIRR ML, accessible both via the command line and via an intuitive Galaxy web interface^47^. immuneML supports the analysis of both BCR and TCR repertoires, with single or paired chains, at the sequence (receptor) and repertoire level. It accepts experimental data in a variety of formats and includes native support for generating synthetic AIRR data to benchmark the performance of AIRR ML approaches. As a flexible platform for tailoring AIRR ML analyses, immuneML features a broad selection of modular software components for data import, feature encoding, ML, and performance assessment (Supplementary Table 1). The platform can be easily extended with new encodings, ML methods, and analytical reports by the research community. immuneML supports all major standards in the AIRR field, uses YAML analysis specification files for transparency, and scales from local machines to the cloud. Extensive documentation for both users and contributors is available (docs.immuneml.uio.no).

immuneML caters to a variety of user groups and usage contexts. The Galaxy web tools make sophisticated ML-based receptor specificity and repertoire immune state prediction accessible to immunologists and clinicians through intuitive, graphical interfaces. The diversity of custom preprocessing and encoding used in published AIRR ML studies hinders their comparison and reproducibility. In contrast, the YAML-based specification of analyses on the command line or through Galaxy improves the collaboration, transparency, and reproducibility of AIRR ML for experienced bioinformaticians and data scientists. The integrated support for AIRR data simulation and systematic ML method benchmarking helps method *users* to select those approaches most appropriate to their analytical setting, and to assists method *developers* to effectively evaluate ML-related methodological ideas.

From a developer perspective, the impressive sophistication of generic ML frameworks such as TensorFlow^68^ and PyTorch^46^ may suggest that these frameworks would suffice as a starting point for AIRR ML method development. However, the fact that the immuneML architecture builds strictly on top of frameworks such as PyTorch underlines the breadth of additional functionality needed for robust ML development and execution in the AIRR domain. For ML researchers, the rich support for integrating novel ML components within existing code for data processing, hyper-parameter optimization, and performance assessment can greatly accelerate method development.

The current version of immuneML includes a set of components mainly focused on supervised ML, but the platform is also suitable for the community to extend it with components for settings such as unsupervised learning^69^ or generative receptor modeling^15,20,70^. We also aim to improve the general support for model introspection, in particular in the direction of supporting causal interpretations for discovering and alleviating technical biases or challenges related to the study design^71^.

In conclusion, immuneML enables the transition of AIRR ML method setup representing a bona fide research project to being at the fingertips of immunologists and clinicians. Complementally, AIRR ML method developers can focus on the implementation of components reflecting their unique research contribution, relying on existing immuneML functionality for the entire remaining computational process. immuneML facilitates the increased adoption of AIRR-based diagnostics and therapeutics discovery by supporting the accelerated development of AIRR ML methods.

## Methods

### immuneML availability

immuneML can be used (i) as a web tool through the Galaxy web interface (galaxy.immuneml.uio.no), (ii) from a command-line interface (CLI), (iii) through Docker (hub.docker.com/repository/docker/milenapavlovic/immuneml), (iv) via cloud services such as Google Cloud (cloud.google.com) through Docker integration, or (v) as a Python library (pypi.org/project/immuneML).

### immuneML analysis specification

immuneML analyses are specified using a YAML specification file (Supplementary Figure 1), which describes analysis components as well as instructions to perform analyses using these components. When using Galaxy, the user may choose to provide a specification file directly or use a graphical interface that compiles the specification for the user. When used as a CLI tool, locally or in the cloud, with or without Docker, the specification file is provided by the user. Examples of specification files and detailed documentation on how to create them are available at docs.immuneml.uio.no/tutorials/how_to_specify_an_analysis_with_yaml.html.

immuneML supports different types of instructions: (i) training and assessment of ML models, (ii) applications of trained ML models, (iii) exploratory data analysis, and (iv) generation of synthetic AIRR datasets. Tutorials detailing these instructions are available at docs.immuneml.uio.no/tutorials.html.

### immuneML public instance

the immuneML Galaxy web interface is available at galaxy.immuneml.uio.no. In addition to core immuneML components, the Galaxy instance includes interfaces towards the VDJdb^56^ database and the iReceptor Gateway^55^. The documentation for the Galaxy immuneML tools is available at docs.immuneml.uio.no/galaxy.html.

### immuneML architecture

immuneML has a modular architecture that can easily be extended (Supplementary Figure 2). In particular, we have implemented glass-box extensibility mechanisms^72^, which enable the creation of customized code to implement new functionalities (encodings, ML methods, reports) that might be needed by the users. Such extensibility mechanisms allow the users to adapt immuneML to their specific cases without the need to understand the complexity of the immuneML code. For tutorials on how to add a new ML method, encoding, or an analysis report, see the developer documentation: docs.immuneml.uio.no/developer_docs.html.

### Use cases

#### Use case 1: Reproduction of a published study inside immuneML

We reproduced the study by Emerson and colleagues using a custom implementation of the encoding and classifier described in the original publication^6^. Out of the 786 subjects listed in the original study, we removed 103 subjects (1 with missing repertoire data, 25 with unknown CMV status, 3 with negative template counts for some of the sequences, and the rest with no template count information, all of which occurred in cohort 1), and performed the analysis on the remaining 683 subjects. We achieved comparable results to the original publication, as shown in Supplementary Figure 4. Supplementary Table 3 shows TCRβ receptor sequences inferred to be CMV-associated, comparing them to those published by Emerson et al.

In addition to reproducing the Emerson et al. study, we retrained the classifier on datasets consisting of 400, 200, 100, and 50 TCRβ repertoires randomly subsampled from cohort 1 and cohort 2. We show how the performance and the overlap of CMV-associated sequences changes with such reductions of dataset size (Figure 2 B and C).

The YAML specification files for this use case are available in the immuneML documentation under use case examples: docs.immuneml.uio.no/usecases/emerson_reproduction.html. The complete collection of results produced by immuneML, as well as the subsampled datasets, can be found in the NIRD research data archive^73^.

#### Use case 2: Extending immuneML with a deep learning component for antigen specificity prediction based on paired-chain (single immune cell) data

To demonstrate the ease of extensibility for the platform, we added a CNN-based receptor specificity prediction ML method to the platform (Supplementary Figure 5), detailing the steps needed to add this as a new component under use case examples in the immuneML documentation: docs.immuneml.uio.no/usecases/extendability_use_case.html. Subsequently, we ran the added component through the standard immuneML model training interface, comparing its predictive performance with TCRdist^17,67^ and logistic regression across three datasets. Additionally, we recovered motifs from the kernels of the neural network by limiting the values of the kernels similar to Ploenzke and Irizarry^74^, and from the hierarchical clustering based on TCRdist distance, and compare these recovered motifs with the motifs extracted by GLIPH2^57^ on the same datasets. Each dataset includes a set of epitope-specific TCRαβ receptors downloaded from VDJdb and a set of naive, randomly paired TCRαβ receptors from the peripheral blood samples of 4 healthy donors^75^. Epitope-specific datasets are specific to cytomegalovirus (KLGGALQAK epitope, with 13000 paired TCRαβ receptors), Influenza A (GILGFVFTL epitope, with 2000 paired TCRαβ receptors), and Epstein-Barr virus (AVFDRKSDAK epitope, with 1700 paired TCRαβ receptors). Dataset details are summarized in Supplementary Table 4. The code for creating the datasets and YAML specifications describing the analysis can be found in the immuneML documentation: docs.immuneml.uio.no/usecases/extendability_use_case.html. The three datasets of epitope-specific receptors, the complete collection of kernel visualizations produced by immuneML, as well as the results produced by GLIPH2, have been stored in the NIRD research data archive^76^.

#### Use case 3: ML methods benchmarking on ground-truth synthetic data

To show immuneML’s utility for benchmarking AIRR ML methods, we constructed a synthetic AIR dataset with known implanted ground-truth signals and performed a benchmarking of ML methods and encodings inside immuneML. To create the dataset for this use case, 2000 human IgH repertoires of 10^5^ CDR3 amino acid sequences were generated using OLGA^62^. Subsequently, immuneML was used to simulate five different immune events of varying complexity by implanting signals containing probabilistic 3-mer motifs (Supplementary Table 2). The signals of each immune event were implanted in 50% of the repertoires, without correlating the occurrence of different immune events. Signals were implanted in 0.1% of the CDRH3 sequences of the repertoires selected for immune event simulation.

Using immuneML, three different ML methods (logistic regression, random forest, support vector machine) combined with two encodings (3-mer and 4-mer frequency encoding) were benchmarked. Hyperparameter optimization was done through nested cross-validation. For the model assessment (outer) cross-validation loop, the 2000 repertoires were randomly split into 70% training and 30% testing data, and this was repeated three times. In the model selection (inner) cross-validation loop, 3-fold cross-validation was used. The test set classification performances of the trained classifiers for each immune event are shown in Figure 2 H.

The immune signals implanted in this dataset can be used to examine the ability of the ML methods to recover ground-truth motifs by comparing the coefficient value (logistic regression, support vector machine) or feature importance (random forest) of a given feature with the overlap between that feature and an implanted signal (Figure 2 I, Supplementary Figure 6).

The bash script for generating the OLGA sequences, as well as the YAML specification files describing the simulation of immune events and benchmarking of ML methods are available in the immuneML documentation under use case examples: docs.immuneml.uio.no/usecases/benchmarking_use_case.html. The benchmarking dataset with simulated immune events as well as the complete collection of figures (for all cross-validation splits, immune events, ML methods, and encodings) can be downloaded from the NIRD research data archive^77^.

## Acknowledgements

We acknowledge generous support by The Leona M. and Harry B. Helmsley Charitable Trust (#2019PG-T1D011, to VG and TMB), UiO World-Leading Research Community (to VG and LMS), UiO:LifeScience Convergence Environment Immunolingo (to VG and GKS), EU Horizon 2020 iReceptorplus (#825821) (to VG), a Research Council of Norway FRIPRO project (#300740, to VG), a Research Council of Norway IKTPLUSS project (#311341, to VG and GKS), the National Institutes of Health (P01 AI042288 and HIRN UG3 DK122638 to TMB) and Stiftelsen Kristian Gerhard Jebsen (K.G. Jebsen Coeliac Disease Research Centre) (to LMS and GKS). We acknowledge support from ELIXIR Norway in recognizing immuneML as a national node service.

## Author contributions

MP, VG, GKS conceived the study. MP and GKS designed the overall software architecture. MP, LS, and KM developed the main platform code. MP and LS performed all analyses. MP, LS, CK, FLMB, RA, GSAH, GB, MC, RF, IG, SG, PHH, KR, ER, PAR, AS, DT, CW, and MW created software or documentation content. RK, NV, KW, LS, MP, AAC, and BC designed and developed the Galaxy tools. CK, RA, TB, MC, SC, LGC, IHH, EH, SH, GK, MLK, CLA, AM, TM, JP, KR, PAR, AR, IS, LMS and GY provided critical feedback. MP, LS, VG, GKS drafted the manuscript. VG and GKS supervised the project. All authors read and approved the final manuscript and are personally accountable for its content.

## Supplementary Figures

**Supplementary Figure 1.**
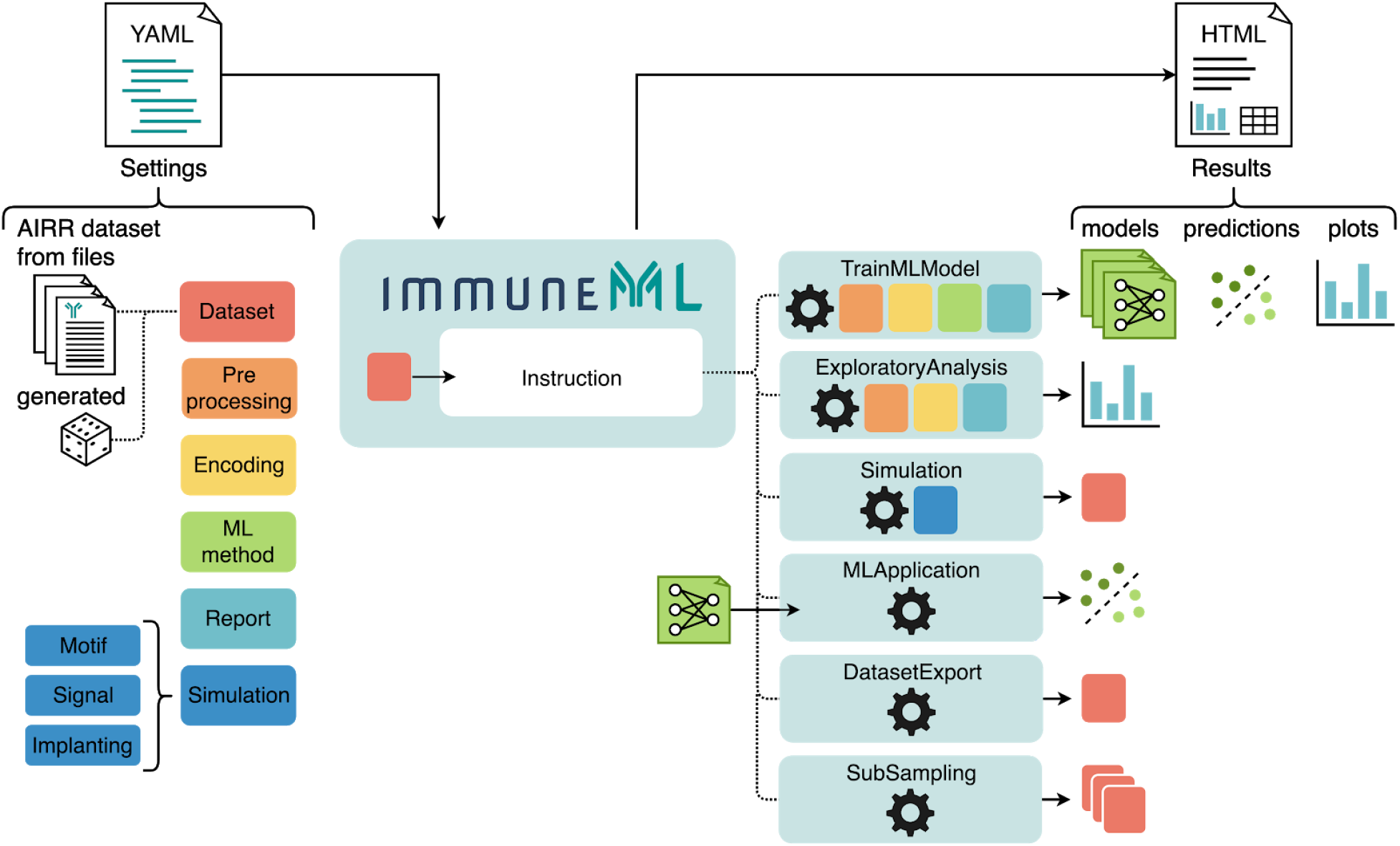
Overview of how immuneML analyses are specified. The YAML specification file describes the analysis components, and what instructions should be performed using these components. The analysis components are datasets, preprocessing, encoding, ML methods, analysis reports, and simulation-related components. Supplementary Table 1 contains a complete list of all components that can be specified. Instructions include training and applying ML models, exploratory analysis, and simulation of synthetic datasets. The results produced by the instructions can be navigated through an HTML summary page generated by immuneML.

**Supplementary Figure 2.**
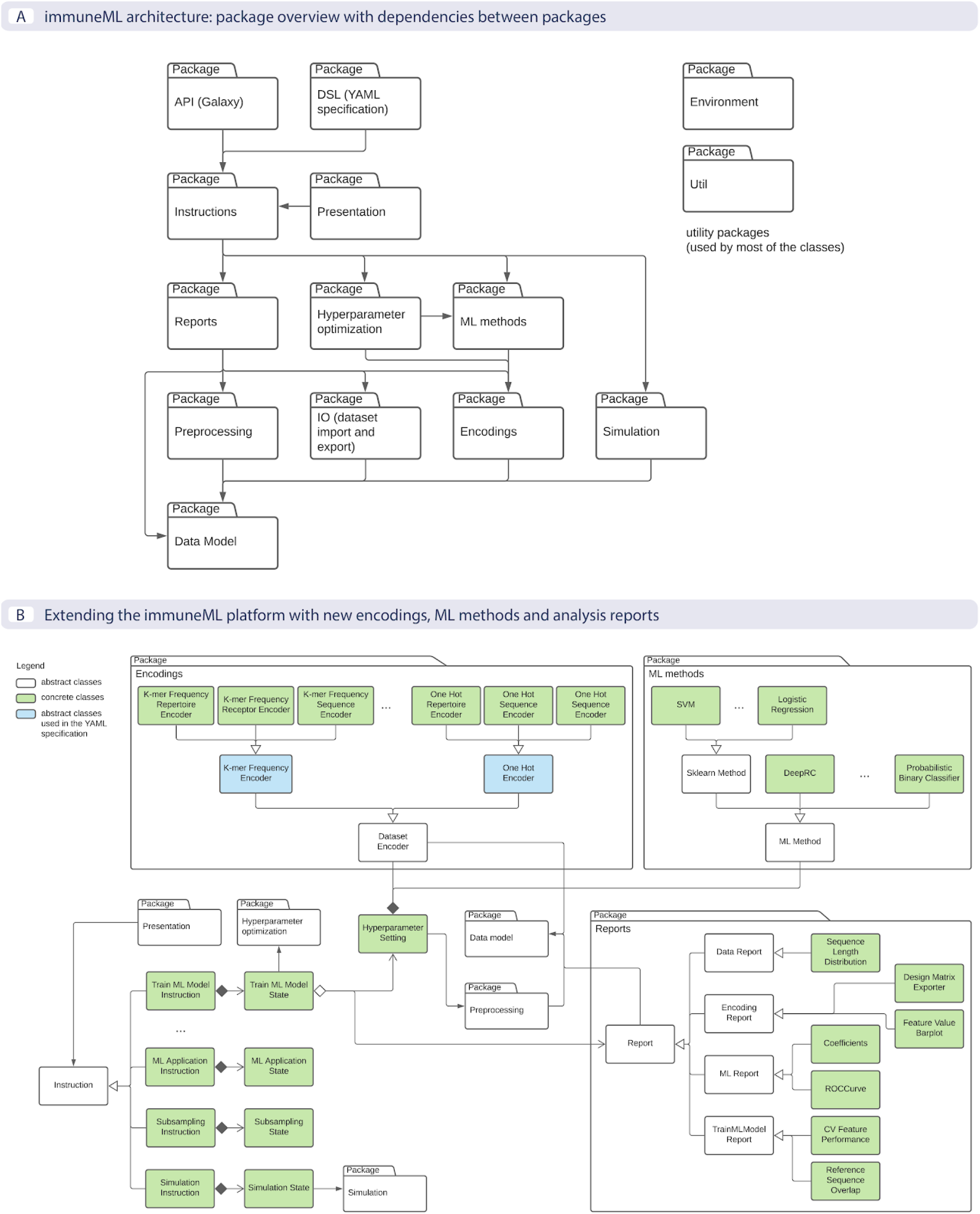
immuneML architecture overview as UML (Unified Modeling Language) diagrams. **A.** High-level overview of the immuneML architecture showing the most important packages and their dependencies. The application programming interface (API) and domain-specific language (DSL) packages represent how the user can interact with immuneML, either through the Galaxy web interface (API package) or when constructing YAML specification files (DSL package). These packages invoke instructions, which map to different analyses that can be performed with immuneML, such as training an ML model or simulating an AIRR dataset. In turn, instructions depend on specific components to perform the analysis. **B.** To extend the platform with new encodings, ML methods, or reports, users may look into the corresponding package and implement the functionalities as described by the appropriate abstract class. The added components could then be used in different instructions according to their purpose. Developer tutorials are available at docs.immuneml.uio.no/developer_documentation.html.

**Supplementary Figure 3.**
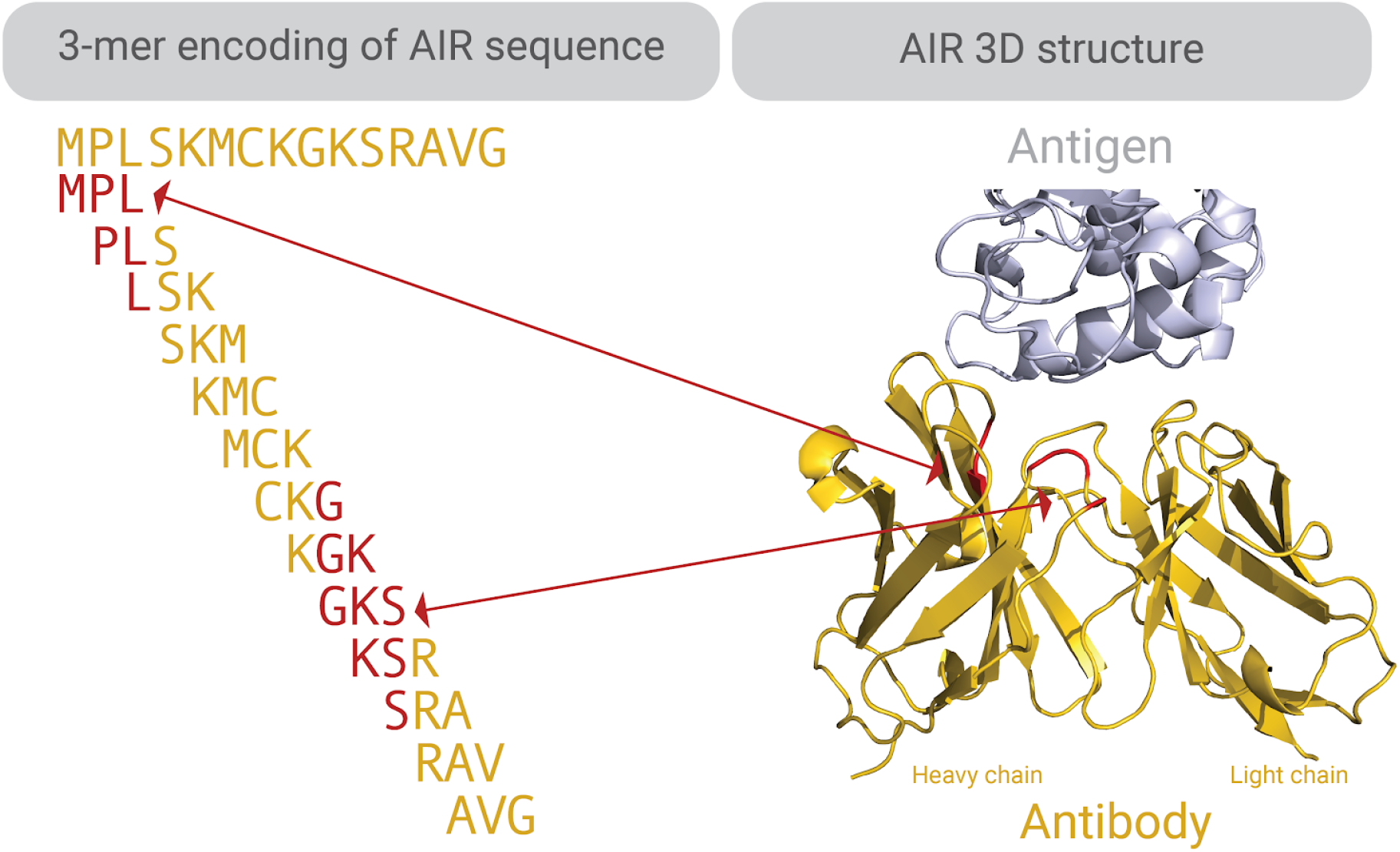
An example of how a k-mer encoded AIR (here antibody) sequence may map the antibody 3D-structure (Protein Data Bank (PDB) ID: 2DQC^78^). K-mers are subsequences of length *k*. Through ML, we can learn, for example, which k-mers are important for determining antigen specificity (color-coded in red). These k-mers may map to regions of the (CDR3) sequence that are in contact with the antigen.^16^ 3D visualization of the antibody-antigen structure was carried out in Pymol^79^.

**Supplementary Figure 4.**
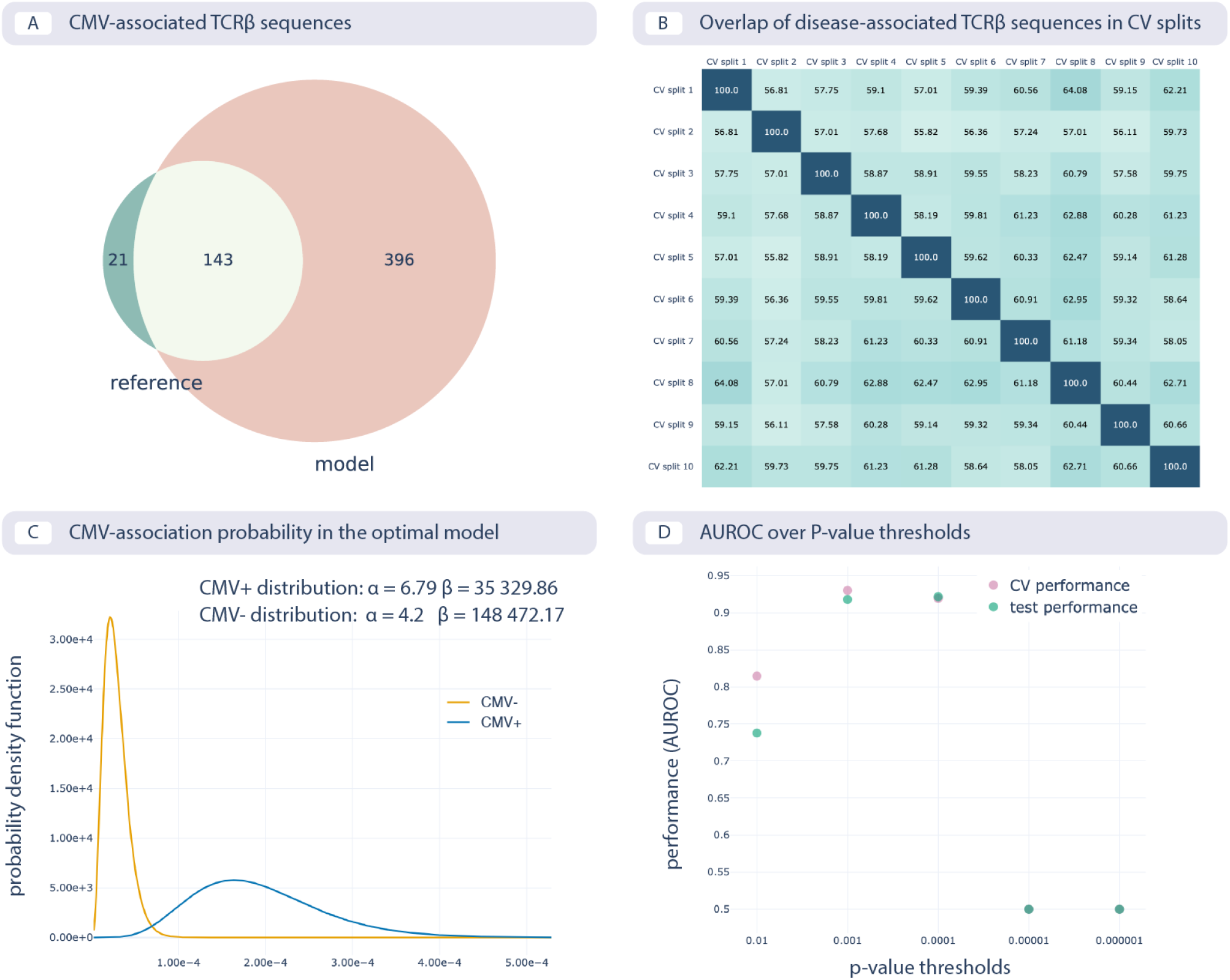
Reproducing the CMV status prediction study by Emerson et al.^6^ **A**. The overlap of the 164 disease-associated TCRβ sequences (V-TCRβaa-J) determined in the original study by Emerson et al., labeled “reference”, with those determined by the optimal model as reproduced here with a p-value threshold of 0.001 (labeled “model”). **B**. The overlap percentage of disease-associated TCRβ sequences for the optimal model with the p-value threshold of 0.001 between different data splits in 10-fold cross-validation (between 50% and 65% overlap). **C.** The probability that a TCRβ sequence is CMV-associated follows a beta distribution estimated separately for CMV positive and negative subjects, which is then used for CMV status prediction of new subjects. **D.** Area under the ROC curve (AUROC) over p-value thresholds in training data (average AUROC over 10 cross-validation splits) and test data (AUROC in cohort 2).

**Supplementary Figure 5.**
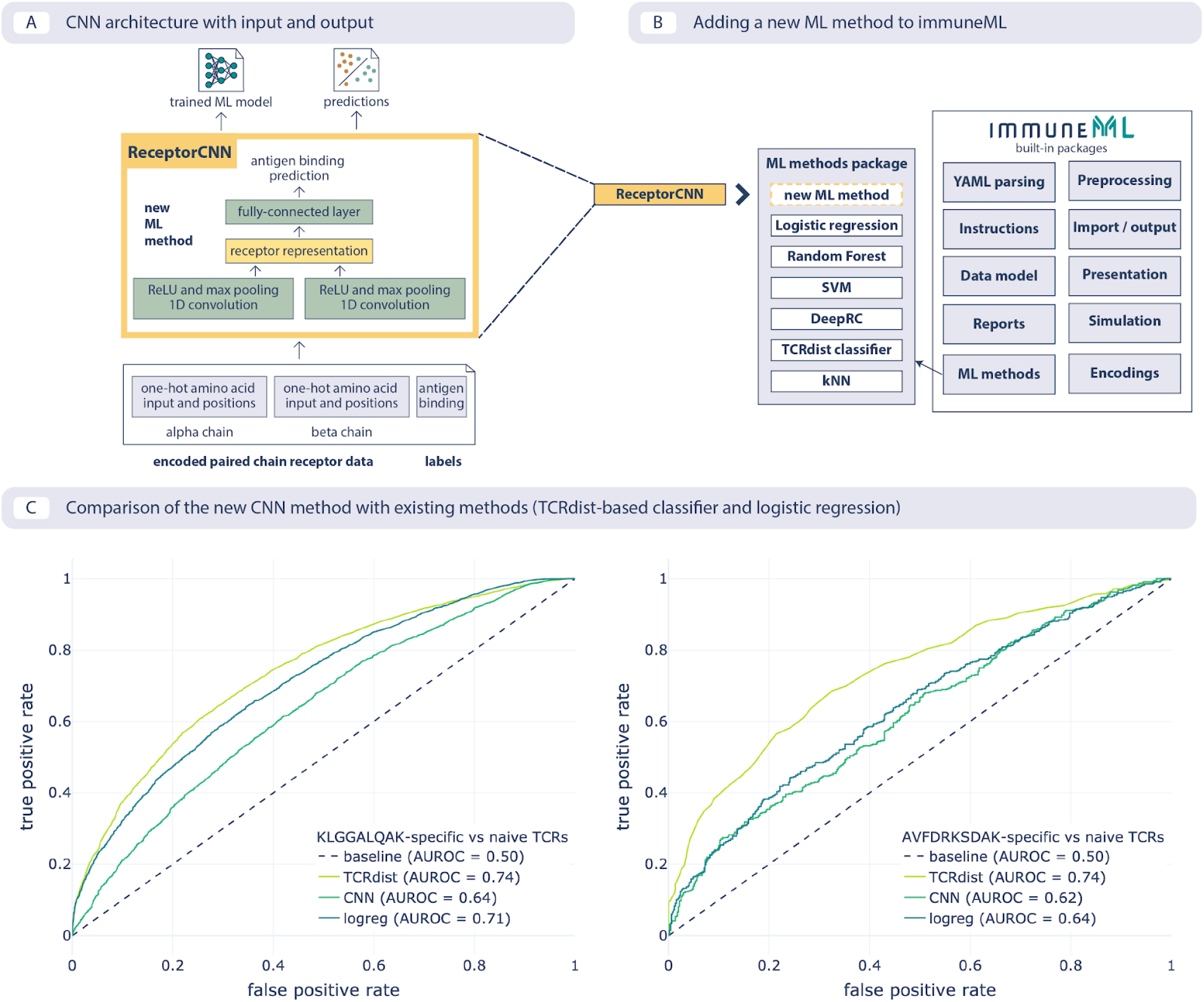
Extending immuneML with a new ML method **A.** The architecture of the new convolutional neural network (CNN) is shown together with expected input in the format of encoded paired-chain receptors and predictions and trained model as output. **B.** Adding a new method requires implementing an interface for the method and then it can reuse the infrastructure (data model, encodings, nested cross-validation, visualizations) without any additional changes. **C**. An example usage where the method can be readily compared with other methods already available within the platform. Here, the area under the ROC curve (AUROC) is shown on two datasets: CMV-specific (epitope: KLGGALQAK, left), and EBV-specific (epitope: AVFDRKSDAK, right), for the CNN that was added (cnn), TCRdist-based k-nearest neighbors classifier (tcrdist) and logistic regression on 3-mer frequencies (logreg).

**Supplementary Figure 6.**
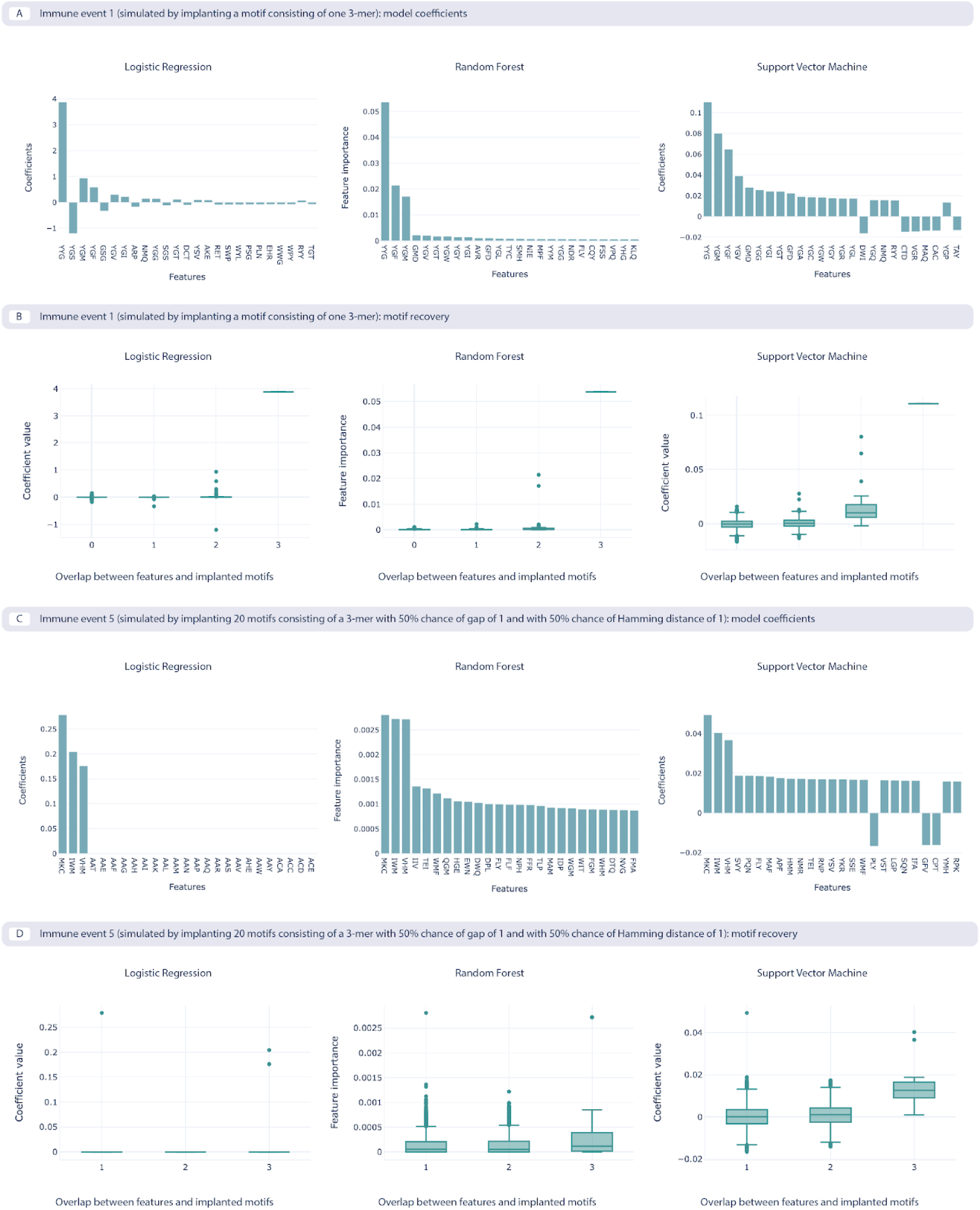
The benchmarking use case model coefficients and motif recovery, where the repertoire data is represented by 3-mer amino acid frequencies. Two immune events are shown. Immune event 1 (A, B) is the simplest event simulated by implanting a single 3-mer, while the immune event 5 (C, D) is the most complex one simulated by implanting 20 motifs consisting of a 3-mer with a 50% chance of having a gap and 50% chance of having a Hamming distance of 1. **A.** The 25 largest coefficients of the logistic regression model, feature importances on random forest model, and coefficients of the support vector machine (SVM) model with a linear kernel. The highest value of the coefficients corresponds to the implanted motif. **B.** Coefficient values for the features depending on the overlap between the recovered features that overlap with the implanted motif, measuring how well the recovered motifs correspond to the implanted motif, shown across the three ML models. **C.** The 25 largest coefficients and feature importances for the ML models trained on immune event 5. **D.** Overlap of recovered and implanted motifs for the ML models trained on immune event 5. Motif recovery for immune event 5 is less effective than for immune event 1.

## Supplementary Tables

**Supplementary Table 1.**
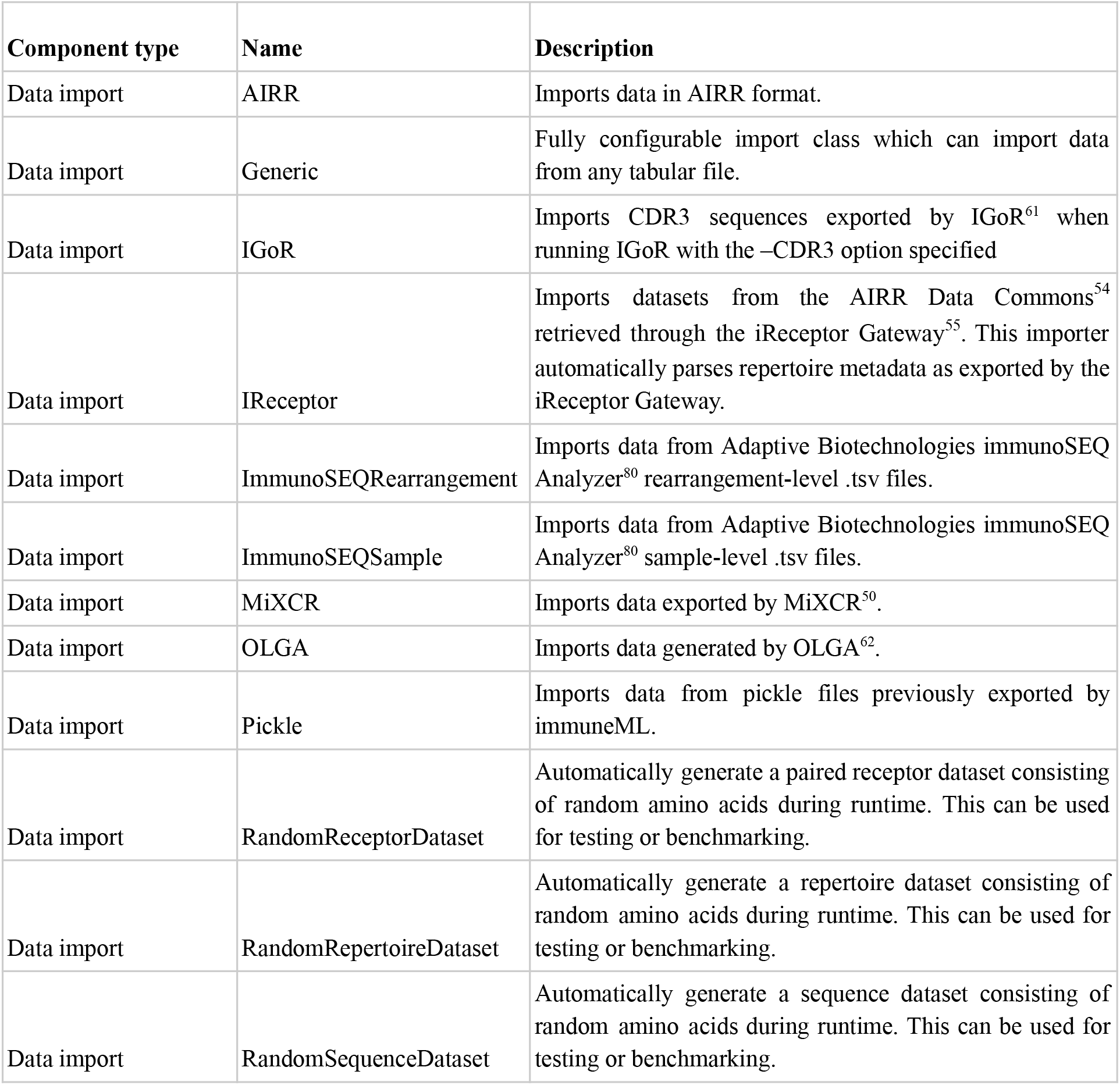
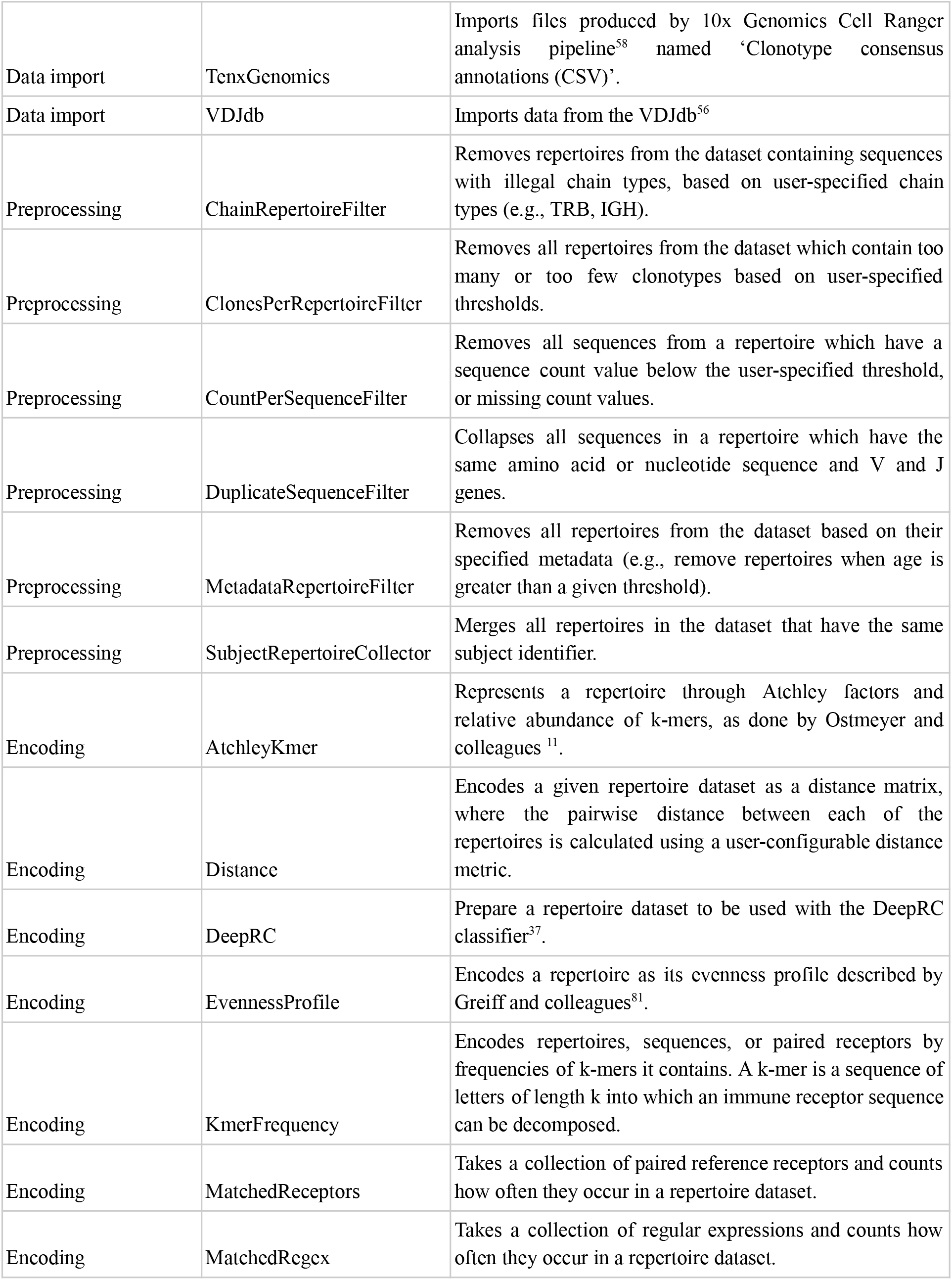
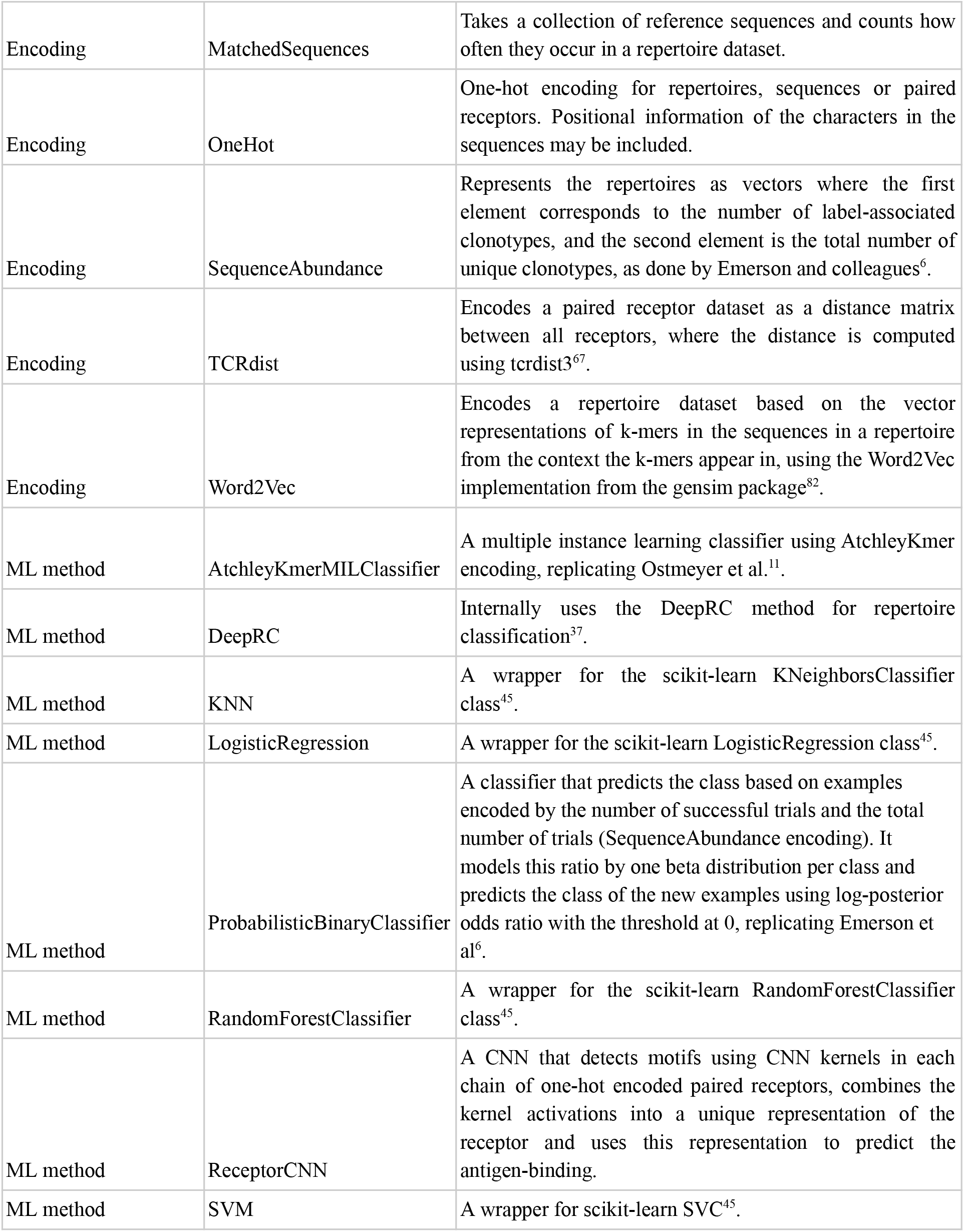
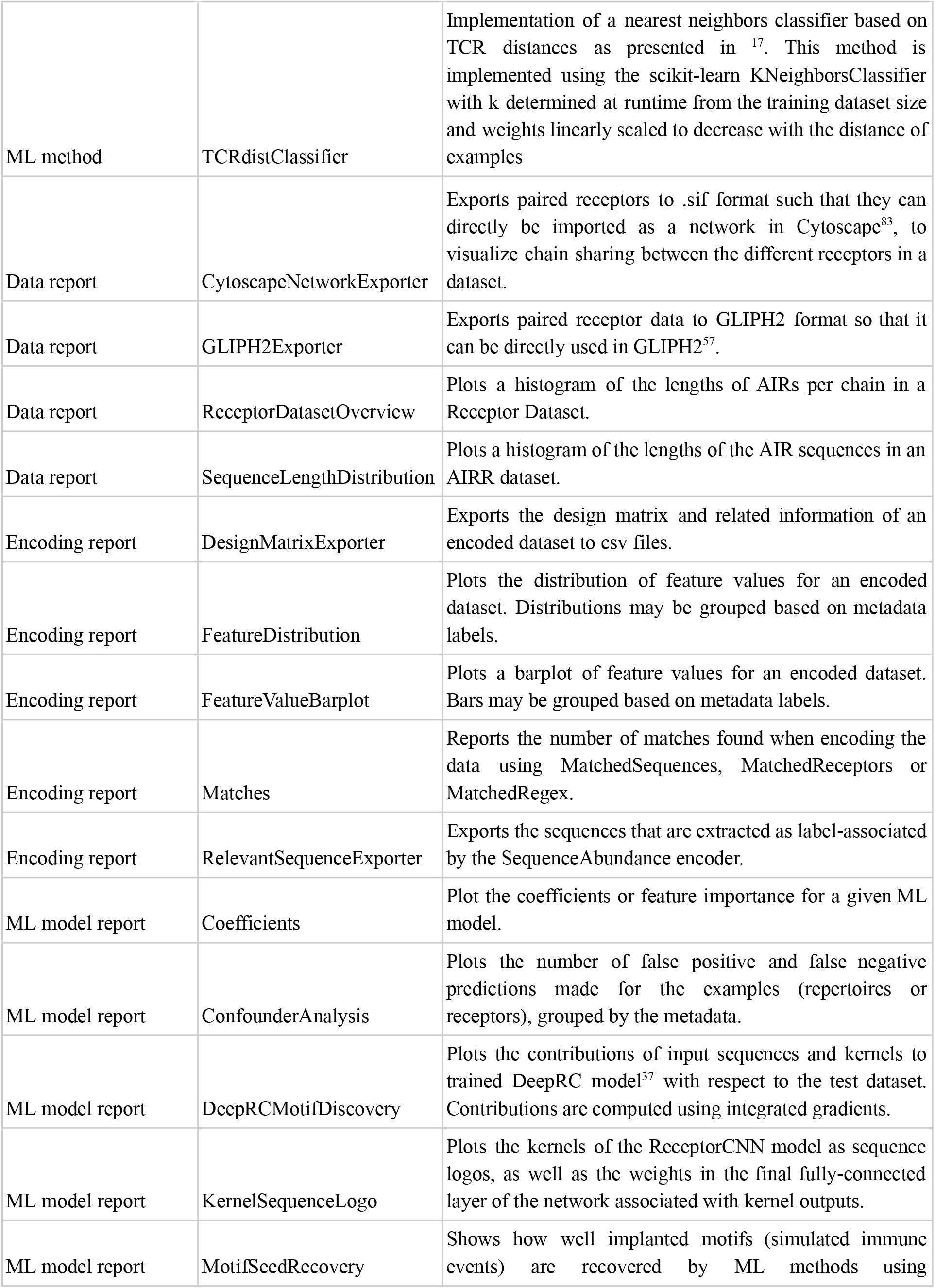
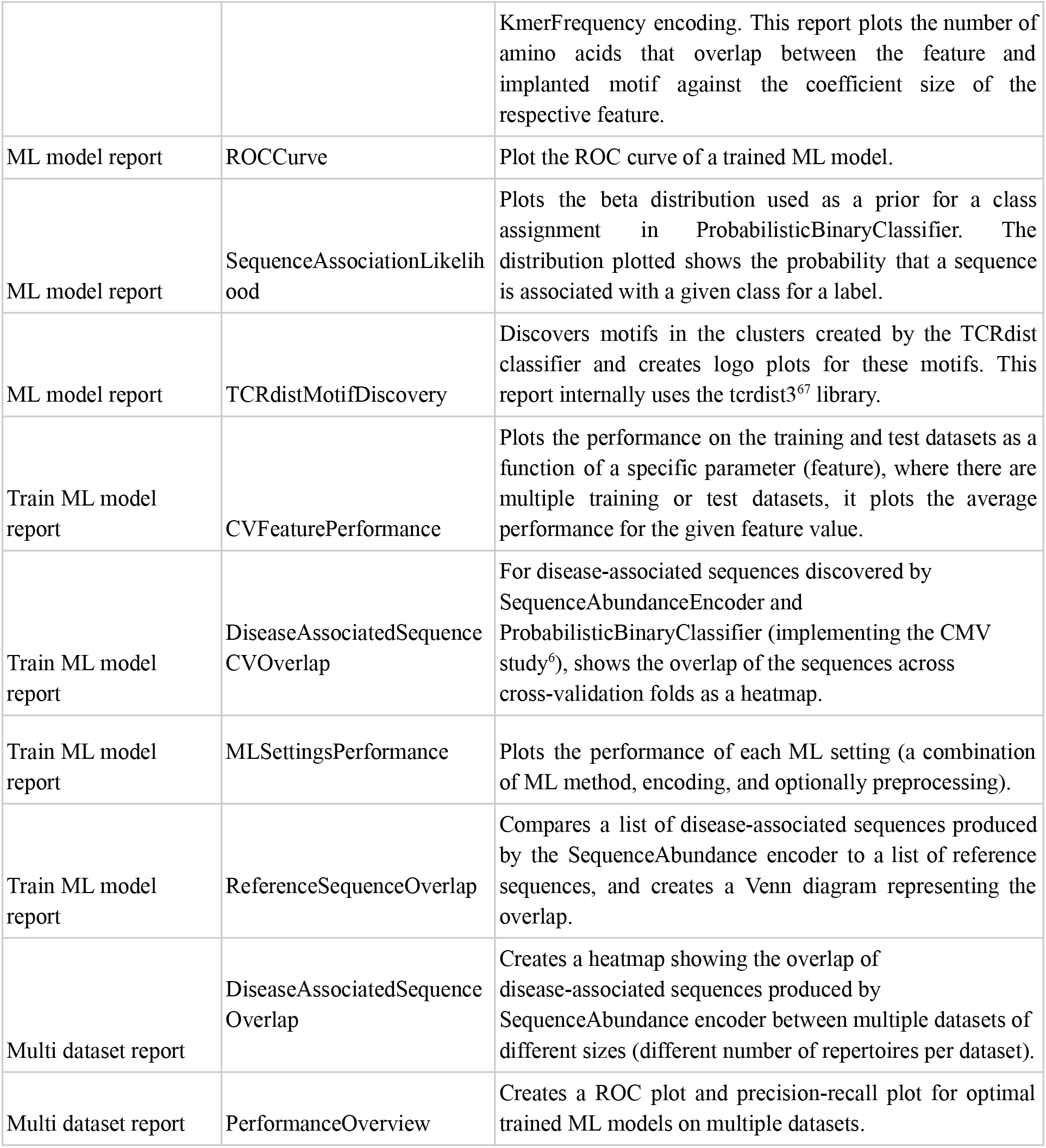
Components for data import, encoding, ML methods, and reports included in immuneML.

**Supplementary Table 2.**
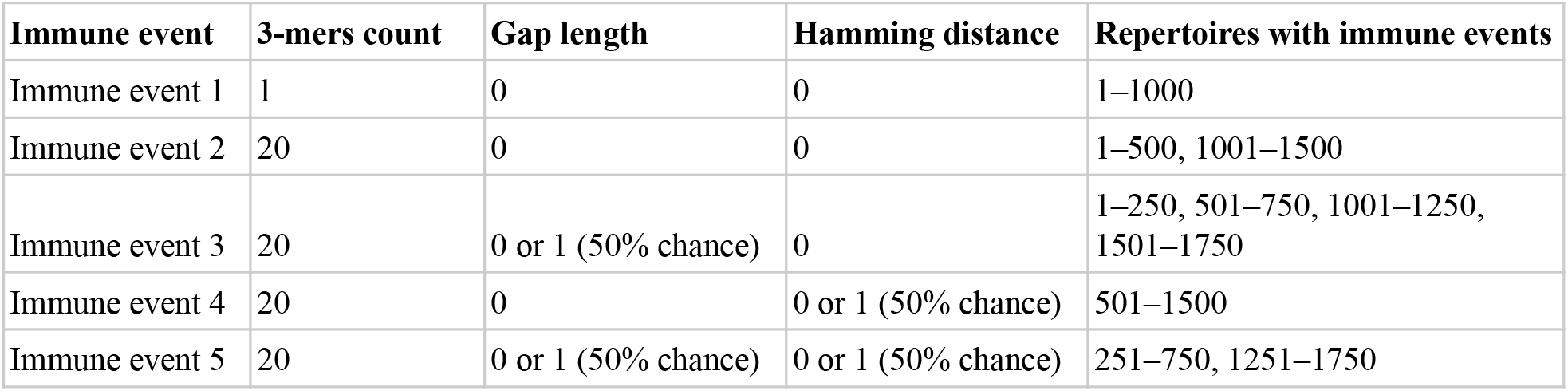
An overview of the settings for the five different simulated immune signals (Figure 2G). One signal can be composed of one or more k-mers (here: 3-mers). For the gapped k-mers, a gap of length 1 was introduced with a 50% chance immediately before or after the middle amino acid of the 3-mer. In the two most complex signals, there was a 50% chance that one of the amino acids in the 3-mer was exchanged for a random different amino acid when implanting it in a sequence. Each immune signal is implanted in a different subset of the repertoires.

**Supplementary Table 3.**
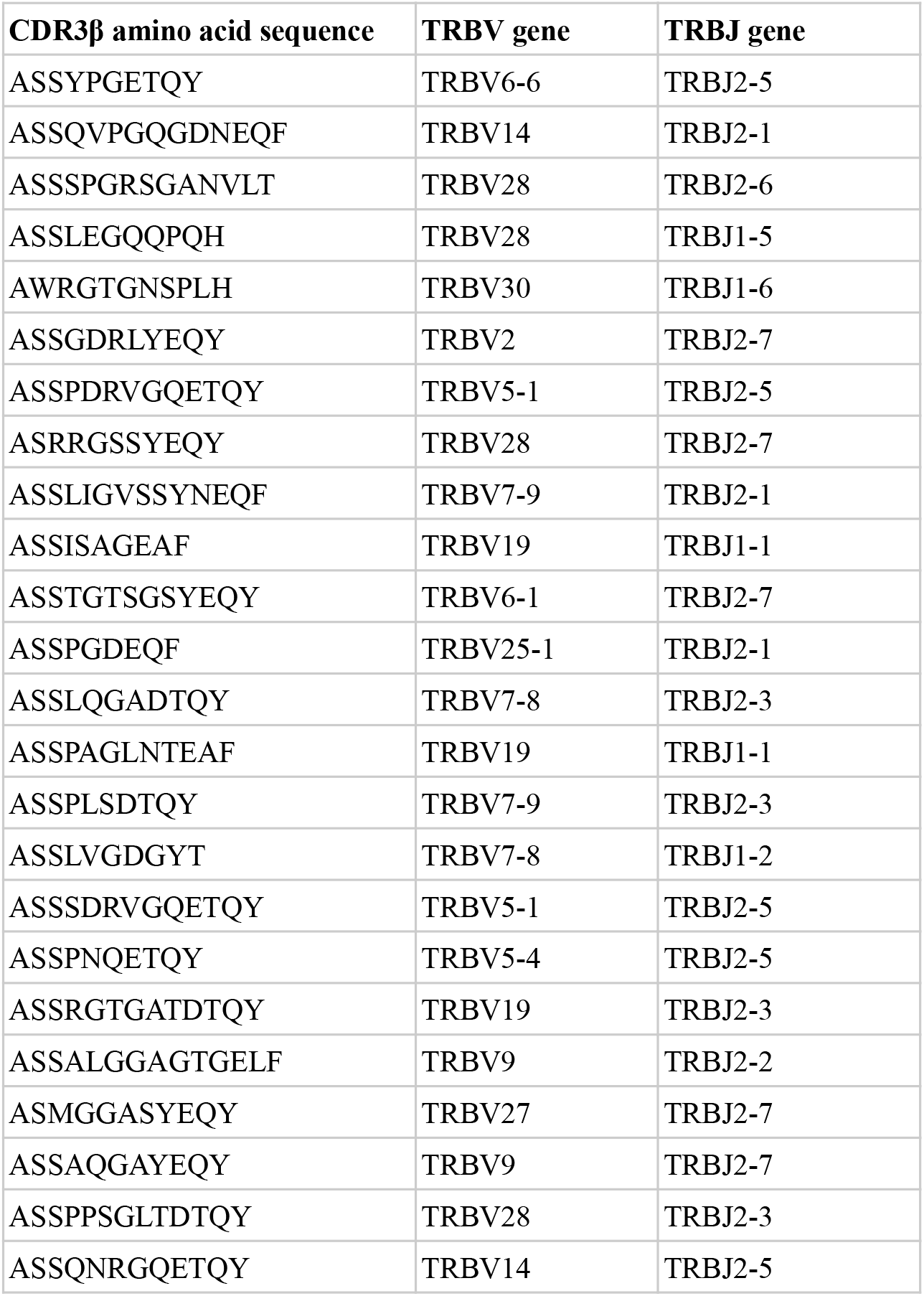
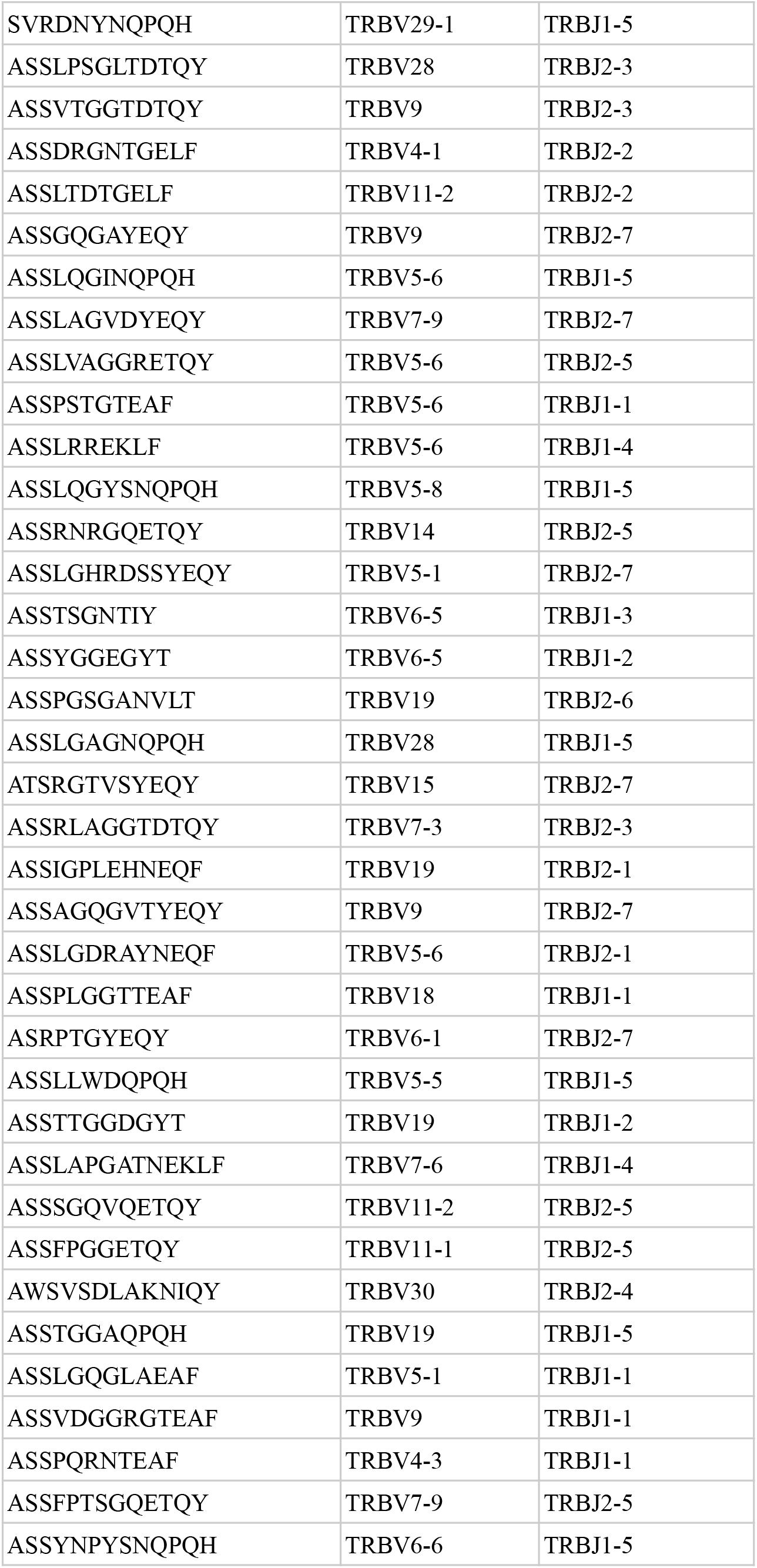
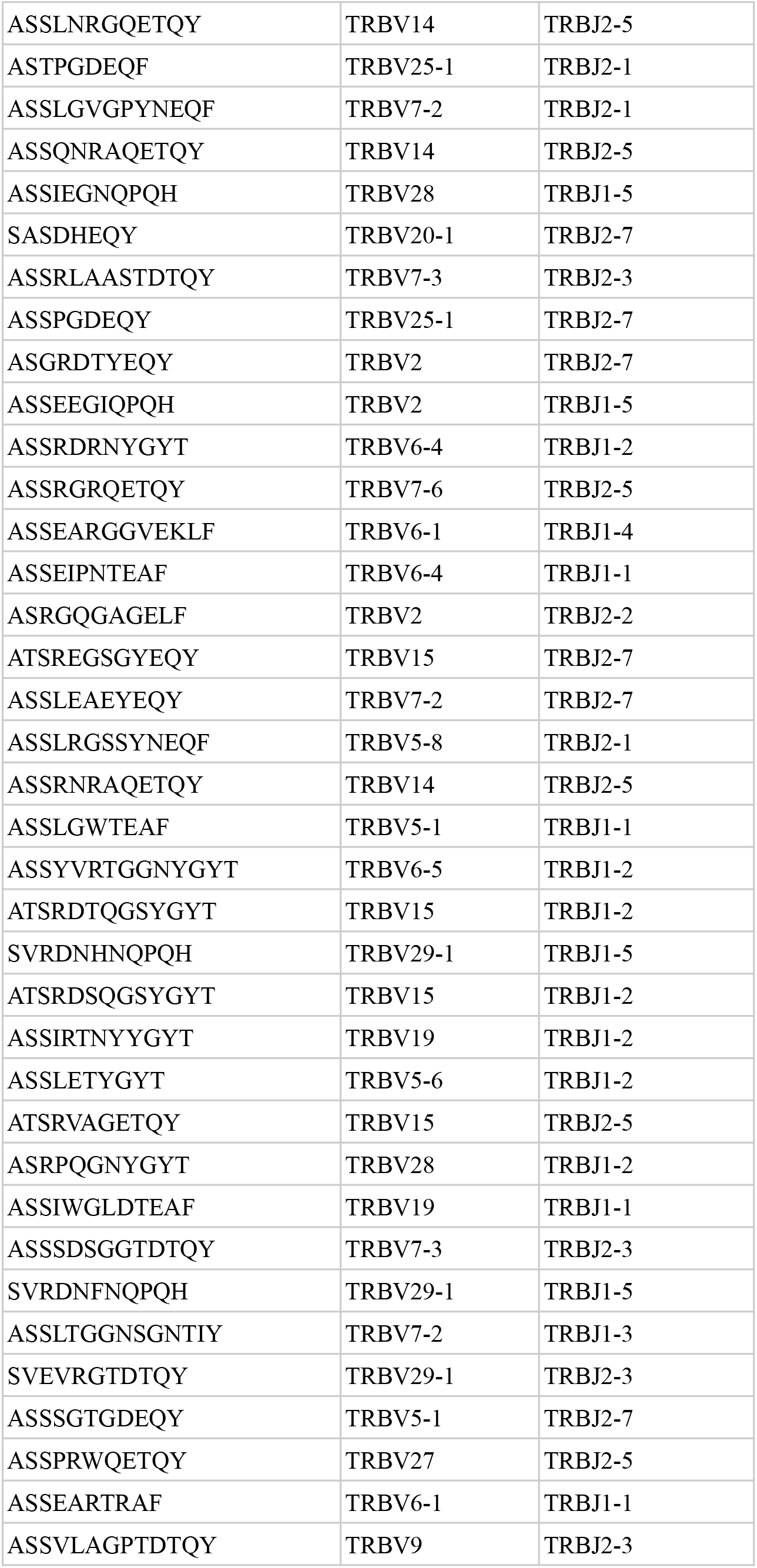
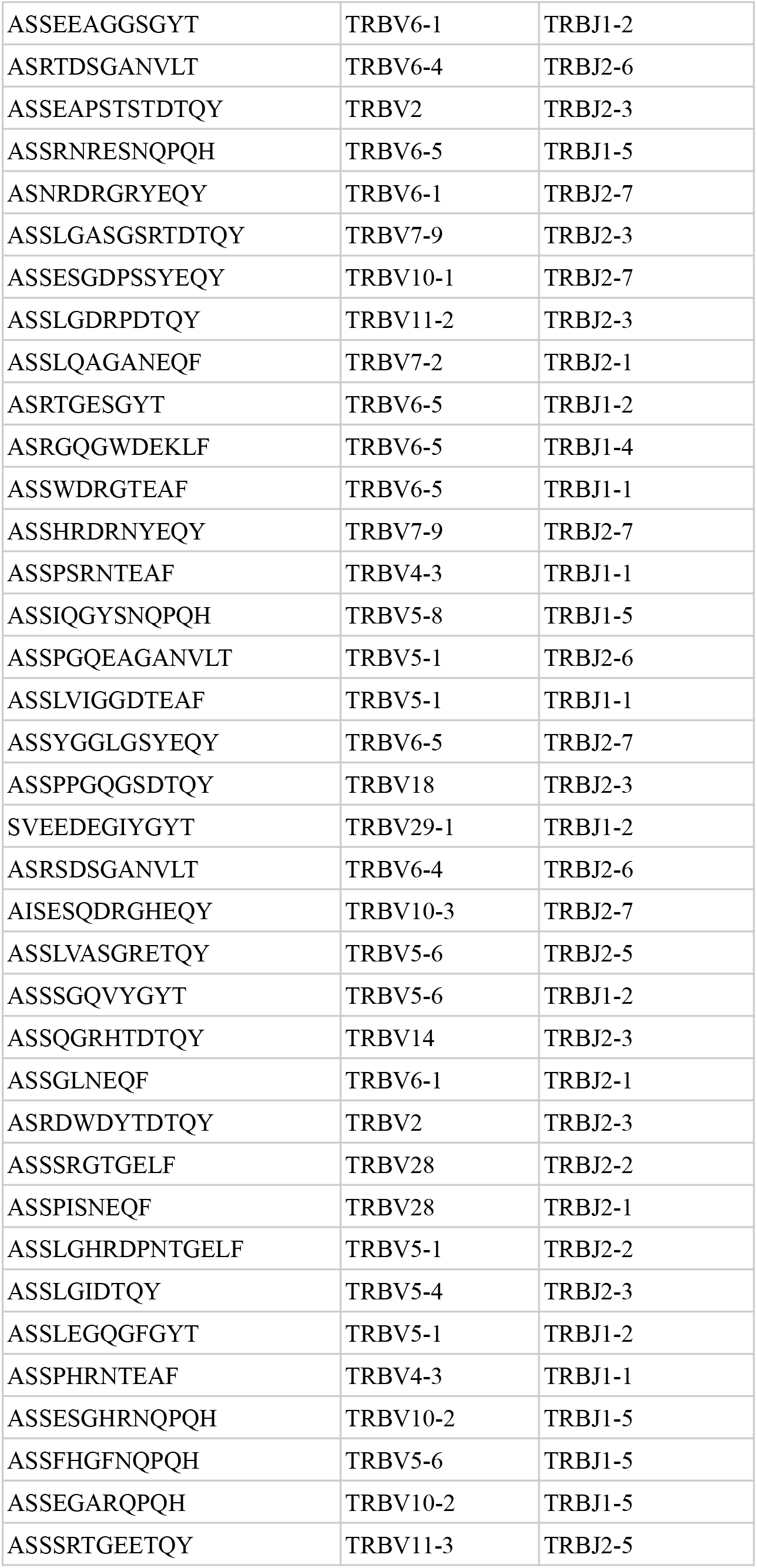
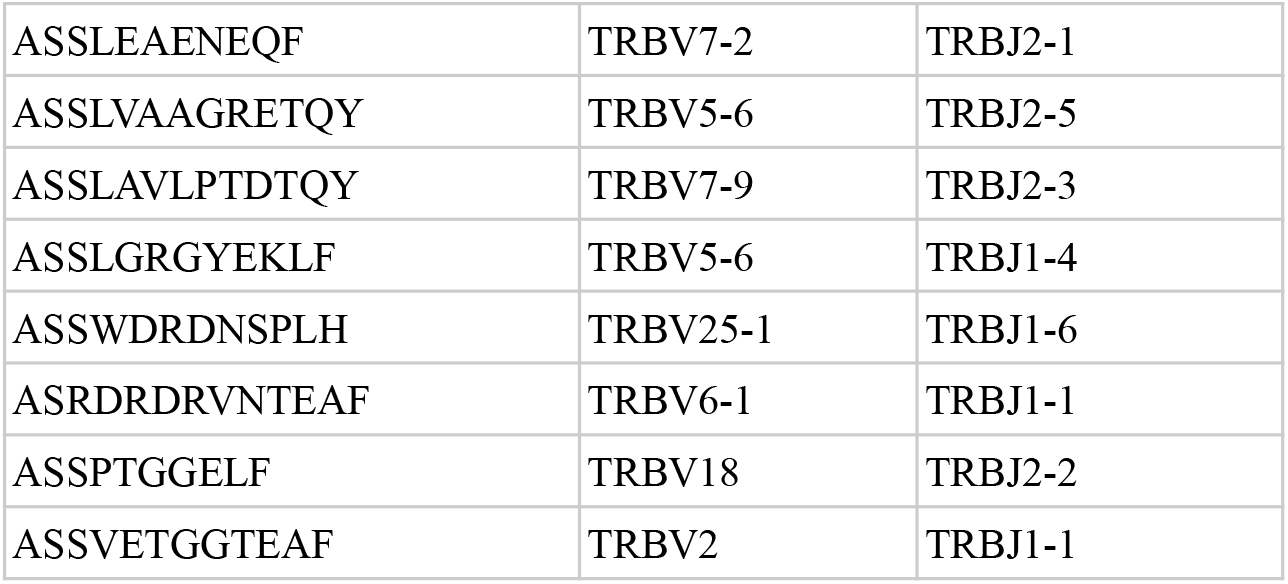
The list of CMV-associated CDR3β sequences with V and J gene determined by training the predictive model on cohort 1 in use case 1 when reproducing the study by Emerson and colleagues^6^ that were also found in the original study. The full list of CMV-associated sequences (including the non-overlapping ones) is available in the NIRD research data archive^73^.

**Supplementary Table 4.** Three epitope-specific datasets in the use case 2.

| Dataset | Number of TCR$\alpha\beta$ receptors | Epitope | Epitope species | Data source: epitope-specific TCRs | Data source: naive receptors | Organism |
| --- | --- | --- | --- | --- | --- | --- |
| AVFDRKSDAK | 3460 | AVFDRKSDAK | EBV | VDJdb<sup>56</sup>: 10x Genomics<sup>84</sup> and EBV study<sup>85</sup> | immuneACCESS: randomly paired TCR$\alpha\beta$ CDR3 from peripheral blood of 4 healthy donors from Heikkilä et al.<sup>75</sup> | Human |
| GILGFVFTL | 4090 | GILGFVFTL | Influenza A | VDJdb<sup>56</sup>: multiple studies<sup>17,18,84,86-90</sup> |  |  |
| KLGGALQAK | 27386 | KLGGALQAK | CMV | VDJdb<sup>56</sup>: 10x Genomics<sup>84</sup> |  |  |

